# Fake IDs? Widespread misannotation of DNA Transposons as a General Transcription Factor

**DOI:** 10.1101/2022.11.24.517799

**Authors:** Nozhat T. Hassan, David L. Adelson

## Abstract

The annotation of transposable elements (TEs) is a critical part of our understanding of genomes, however, the accuracy of annotation pipelines remains an issue as TEs are frequently underestimated or misannotated. We report the General Transcription Factor II-I Repeat Domain-Containing Protein 2 (GTF2IRD2) was used to erroneously annotate DNA transposons in a variety of non-mammalian species as GTF2IRD2 contains a 3’ fused hAT transposase domain. This study emphasises that the misannotation of TEs as *trans-*regulatory elements, such as transcription factors, can lead to errors in phylogenetic trees based on orthologs and to significant wasted time for investigators interested in gene regulation and characterising non-mammalian genomes.

## Background

Annotation of TEs is essential for understanding genome structure and function; however, misannotation may result in erroneous classification of TEs as *trans-*regulatory elements such as transcription factors. TEs can introduce genetic novelty to the host genome and an example of this is the exaptation of TEs into the human General Transcription Factor II-I Repeat Domain-Containing Protein 2 (GTF2IRD2). GTF2IRD2 contains a Charlie8-like element positioned at the 3’ end (C terminus) of the gene model/full protein that has retained transposable element features such as DDE amino acids required for transposition [1]. The Charlie transposon (DNA transposon; hAT superfamily) is an old autonomous group of transposons abundant in mammalian genomes, including humans [2]. Charlie transposons are defined by their target site duplication (TSDs) and terminally inverted repeats (TIRs). At the same time, the protein-coding sequence of the transposase may vary between Charlie elements from different species [3]. GTF2IRD2 is found in mammals and is predicted to be in many reptiles, amphibians and bony fishes [4]. However, upon closer inspection, some non-mammalian GTF2IRD2 sequences appear to be hAT transposons, not transcription factors. Here we have used structural and phylogenetic analysis to resolve the widespread misannotation of non-mammalian DNA transposons as GTF2IRD2 transcription factors. We believe that this paper addresses an important issue; it has implications not only for the study of TEs but is also relevant as similar misannotations could also cause the misinterpretation of other results that depend on reliable gene annotation.

## Results and Discussion

While annotating hAT-6 transposons in Testudines genomes, we noticed that GTF2IRD2/2A, a human general transcription factor was the top BLASTN result when using hAT-6 transposons as a query to search non-mammalian genomes. This was unexpected as hAT-6 has hallmarks of a functional transposase, such as TIRs and TSDs, and has the functional motifs required for transposition, such as the DDE/RW residues in the translated open reading frame (ORF) [4,5]. NCBI’s eukaryotic gene annotation pipeline prefers to use experimental evidence when annotating genes but uses an *ab initio* model to predict optimal CDS alignments when there is no experimental data (https://www.ncbi.nlm.nih.gov/genome/annotation_euk/process/). Specifically, Gnomon is meant to exclude gene predictions with high homology to transposable or retro-transposable elements from the final gene models, however, the eukaryotic annotation pipeline appears to lack a final TE filtering step after integrating RefSeq annotations. This may explain how genes such as GTF2IRD2/2A, which contains an integrated Charlie8-like element (a hAT-like transposase), can lead to hAT-6 being predicted to be a transcription factor in non-mammalian genomes [3]. When searching Interpro for GTF2IRD2/2A sequences, we saw that 485 proteins were annotated as GTF2IRD2/2A proteins, but only 4 of these have been reviewed in human, cow and mouse genomes (https://www.ebi.ac.uk/interpro/entry/InterPro/IPR042224/protein/reviewed/#table, Accessed 11th November 2022). To determine if additional hAT transposons were incorrectly annotated as GTF2IRD2/2A, we examined the phylogeny of a set of protein sequences annotated as GTF2IRD2/2A or GTF2IRD2/2A-like from mammals, reptiles, bony fishes, and amphibians (fig. 1). Birds were excluded from analysis as the sequences annotated as GTF2IRD2/2A had no significant similarity to either mammalian GTF2IRD2/2A or to any DNA TE structures. Any sequence homology in birds was limited to the N terminus of the mammalian GTF2IRD2/2A, an indication of potential similarity to the ancestral GTF2IRD2/2A protein prior to the exaptation of Charlie8 (Supplementary Materials).

**Figure 1:**
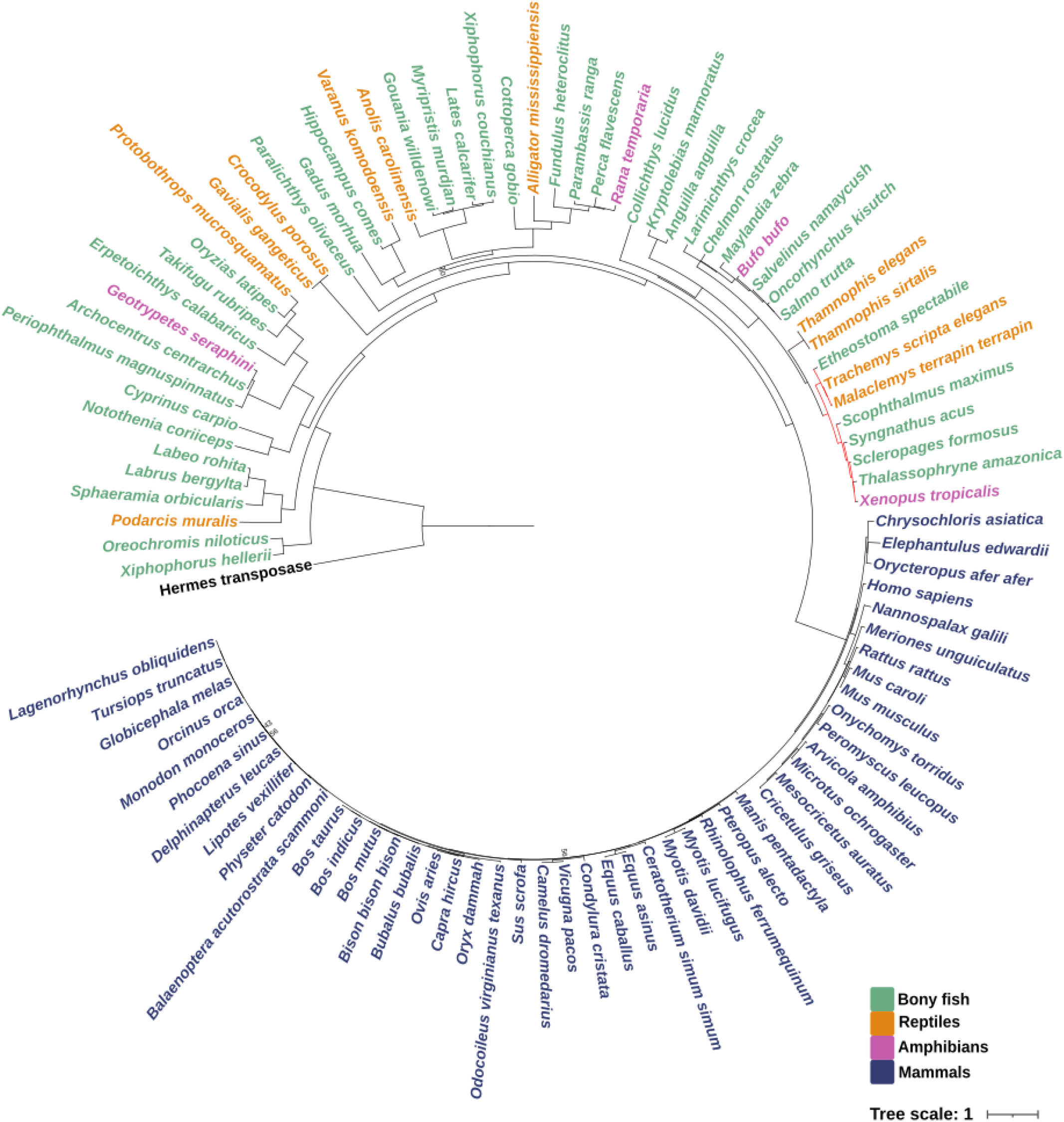
The phylogenetic relationship of sequences annotated as GTF2IRD2/2A from bony fishes, reptilian, amphibian, and mammalian genomes from NCBI. Bony fish are coloured green, reptiles are orange, amphibians are pink and mammals are blue. Branches labelled in red are hAT-6 transposons. The Hermes transposase was used as an outgroup. Support values under 60 are shown at nodes. For the full tree and support values, see Supplementary materials.

There was a clear distinction between mammalian and non-mammalian GTF2IRD2/2A gene models. Mammalian GTF2IRD2/2A are correctly annotated as transcription factors as they have an N terminus ∼400 aa long containing a GTF2I-Like repeat domain, a zinc finger binding domain, and an integrated Charlie8-like element at the C terminus [1]. However, this is not the case with non-mammalian sequences. Multiple alignment of GTF2IRD2/2A from mammals and non-mammals with hAT-6 transposons exclusively shows high similarity alignment at the 3’ end (fig 2). This is consistent with the position of the Charlie-like element in GTF2IRD2/2A, demonstrating that non-mammalian GTF2IRD2/2A are not likely to be TFs, but TEs.

**Figure 2:**
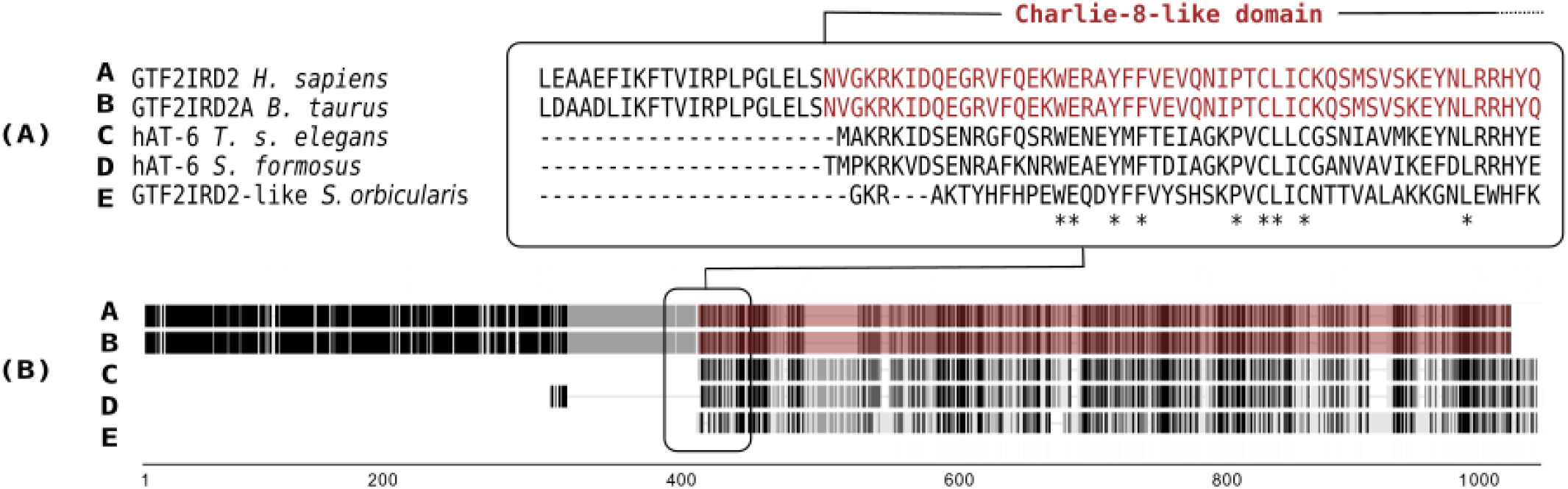
(A) Boundary of non-transposon and Charlie8-like domain of a multiple sequence alignment (MSA) of mammalian GTF2IRD2/2A to hAT-6 transposons and a TE sequence misannotated as GTF2IRD2-like. (B) Full schematic MSA of the selected sequences. The length of sequences in amino acids is shown on the x-axis from the 5’ to 3’ direction. The Charlie8-like domain of mammalian GTF2IRD2/2A is highlighted in red text and red shading.

Non-mammalian GTF2IRD2/2A from fish, reptiles, and frogs were manually curated to identify signature motifs of DNA transposons as they appeared closely related to hAT-6 transposons than mammalian GTF2IRD2/2A. 28 sequences were classified as autonomous hAT transposons from several different species, but particularly from the genome of the Atlantic salmon (*Salmo salar*) (Table 1). TSDs and TIRs are characteristic of hAT transposons and were found in most sequences, while sequences without them were classified as partial, non-autonomous transposons. Furthermore, DDE/RW residues required for transposition were identified for most of the newly curated hAT transposons (Supplementary Materials). Finally, the 5’ and 3’ TIRs were mostly conserved across the misannotated hAT transposons, which further demonstrates the degree of misannotation (fig. 3).

**Table 1:** Species from the *Actinopterygii, Reptilia* and *Amphibia* class containing hAT transposons that were incorrectly annotated as GTF2IRD2/2A. The best match of each hAT transposon to the Repbase database and the corresponding pairwise identity is shown.

| Class | Species Name | Length (bp) | Repbse Best Match | Pairwise Identity (%) |
| --- | --- | --- | --- | --- |
| *Actinopterygii* | *Sphaeramia orbicularis* | 2921 | hAT-1_SSa | 81 |
|  | *Xiphophorus hellerii* | 3231 | hAT-1_XT | 79 |
|  | *Archocentrus centrarchus* | 2522 | hAT-2_PM | 83 |
|  | *Takifugu rubripes* | 2612 | hAT-7_PM | 82 |
|  | *Erpetoichthys calabaricus* | 2591 | hAT-4_SSa | 88 |
|  | *Oryzias latipes* | 2614 | hAT-1_SSa | 67 |
|  | *Xiphophorus couchianus* | 2756 | hAT-1_SSa | 75 |
|  | *Gouania willdenowi* | 2789 | CHAPLIN4_FR | 91 |
|  | *Cottoperca gobio* | 2656 | hAT-1_SSa | 74 |
|  | *Kryptolebias marmoratus* | 2673 | hAT-1_SSa | 74 |
|  | *Larimichthys crocea* | 2787 | hAT-6_XT | 87 |
|  | *Chelmon rostratus* | 2530 | hAT-1_SSa | 80 |
|  | *Maylandia zebra* | 2551 | hAT-1_SSa | 87 |
|  | *Salvelinus namaycush* | 2488 | hAT-1_SSa | 99 |
|  | *Oncorhynchus kisutch* | 2508 | hAT-1_SSa | 98 |
|  | *Salmo trutta* | 2893 | hAT-1_SSa | 99 |
|  | *Labrus bergylta* | 2191 | hAT-2B_PM | 96 |
|  | *Labeo rohita* | 2528 | hAT-8_PM | 72 |
|  | *Paralichthys olivaceus* | 2255 | hAT-17_SSa | 79 |
|  | *Gadus morhua* | 2803 | hAT-22_DR | 70 |
|  | *Lates calcarifer* | 2788 | hAT-9_DR | 83 |
|  | *Collichthys lucidus* | 3127 | CHAPLIN8_FR | 91 |
|  | *Anguilla anguilla* | 3562 | hAT-1_SSa | 95 |
|  | *Perca flavescens* | 2442 | hAT-1_SSa | 80 |
|  | *Parambassis ranga* | 2630 | hAT-4_SSa | 70 |
| *Amphibia* | *Bufo bufo* | 2392 | hAT-1_SSa | 93 |
| *Reptilia* | *Anolis carolinensis* | 2250 | hAT-2_AC | 100 |
|  | *Alligator mississippiensis* | 4120 | hAT-7_AMi | 87 |

**Figure 3:**
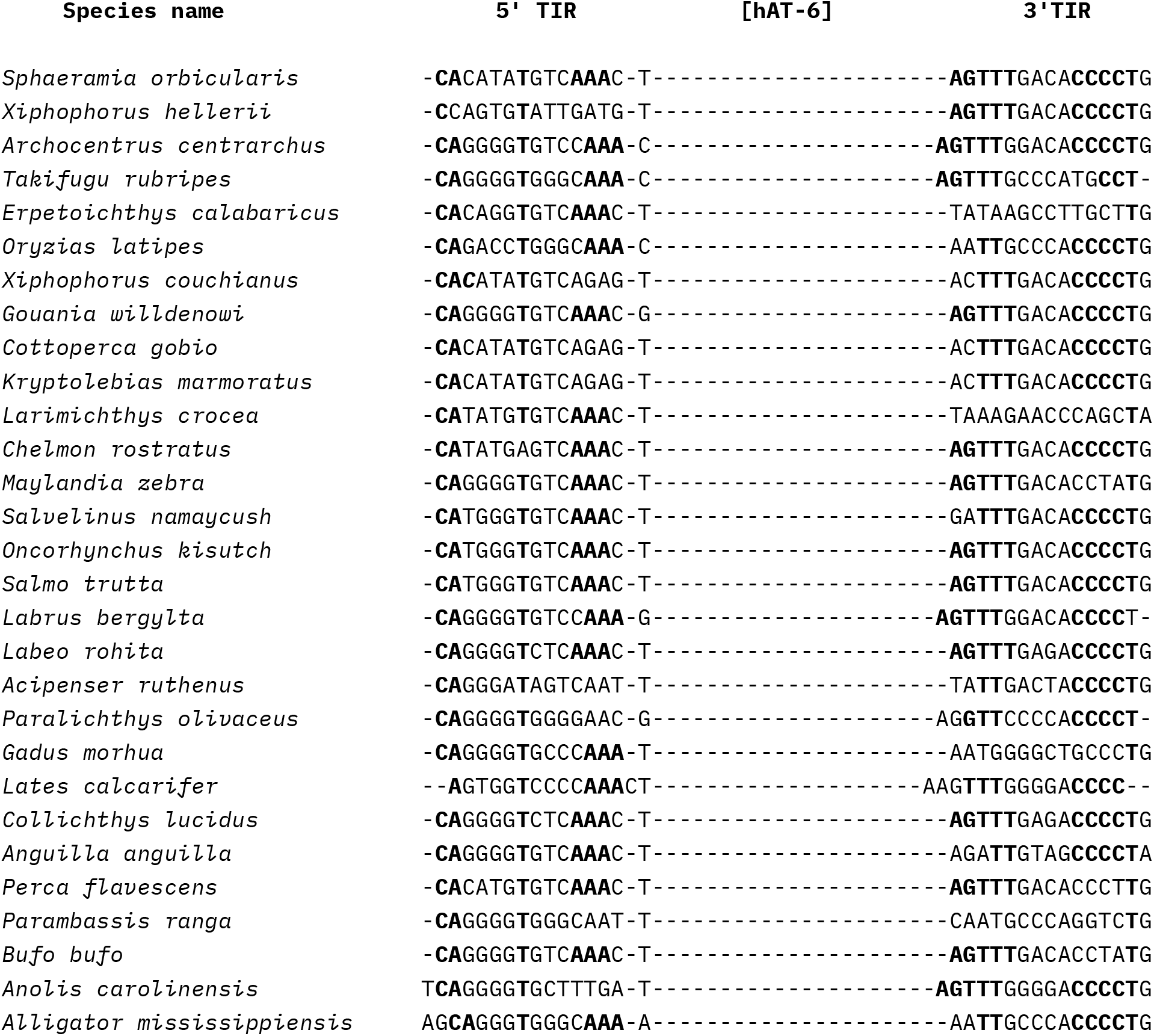
Multiple alignment of the 5’ and 3’ TIRs of *Actinopterygii, Amphibia* and *Reptilia* hAT transposons derived from sequences misannotated as GTF2IRD2/2A. Nucleotides in bold text are conserved in most of the species.

## Conclusions

In this study we present a case of widespread incorrect annotation of hAT DNA transposons as GTF2IRD2/2A. This has led to at least 28 or more new instances of hATs from reptiles, bony fishes, and amphibians that had been overlooked and could have been incorrectly used as transcription factors in other analyses. Correct annotation is a vital step in furthering our understanding of genome evolution, and misannotation of TEs as *trans-*regulatory genes such as TFs affects downstream research and can confound phylogenetic analysis.

## Methods

### Manual curation of hAT transposons from GTF2IRD2/2A sequences

A set of GTF2IRD2/2A and GTF2IRD2/2A-like protein sequences were downloaded from NCBI. The search was limited to species belonging to the *Actinopterygii, Reptilia* and *Amphibia* classes. To determine whether these GTF2IRD2/2A proteins were actually DNA transposons, extensive manual curation was performed to locate characteristic sequence features such as TIRs, TSDs and ORFs. To identify hAT transposons, hAT-6_TSE (in prep) was used as a query in a BLASTP search against a set of mammal, reptile, amphibian, bony fish and bird genomes containing GTF2IRD2/2A gene annotations [6]. The corresponding nucleotide sequence of each top hit was extended 1000 bp in flanking regions where possible and used for manual annotation of TIRs and TSDs characteristic of hAT transposons. ORFs were searched using GENSCAN and searched for DD/E and RW residues. Sequences that contained 5’ and 3’ TSDs, TIRs, and an intact ORF were classified as autonomous hAT transposons [6,7] (Supplementary Materials). The best match for each new autonomous hAT was found using Repbase and both 5’ and 3’ TIRs were aligned using MAFFT v7.310 to view conserved nucleotides [8]. hATs misannotated as GTF2IRD2-like were aligned using MAFFT to mammalian GTF2IRD2/2A to confirm they had homology to GTF2IRD2/2A’s Charlie8-like domain (Supplementary Materials).

### Tree-building

GTF2IRD2/2A and GTF2IRD2/2A-like protein sequences from mammals, reptiles, amphibians, and bony fishes were aligned to the Hermes and hAT-6 transposons using MAFFT v7.319 [9]. The alignment was trimmed using CLipKit and IQTree was used for tree reconstruction with JTT+F+I+G4 as the best-fit model with 20 maximum likelihood trees and 1000 bootstraps [10–13].

## Abbreviations

GTF2IRD2: General Transcription Factor II-I Repeat Domain-Containing Protein 2
GTF2IRD2A: General Transcription Factor II-I Repeat Domain-Containing Protein 2A

## Acknowledgements

We would like to thank Terry Bertozzi and Christine Elsik for their expertise and helpful comments. We would also like to thank our lab members for their ongoing support throughout this study and beyond.

## Declarations

### Ethics approval and consent to participate

Not applicable

### Availability of data and materials

The dataset(s) supporting the conclusions of this article are included within the article (and its additional file(s)).

### Authors’ contributions

NTH performed the data processing and analysis. NTH and DLA wrote the manuscript. The authors read and approved the final manuscript

