## Supplementary figures and images for "Fake IDs? Widespread misannotation of DNA Transposons as a General Transcription Factor"

### support_value_tree.pdf

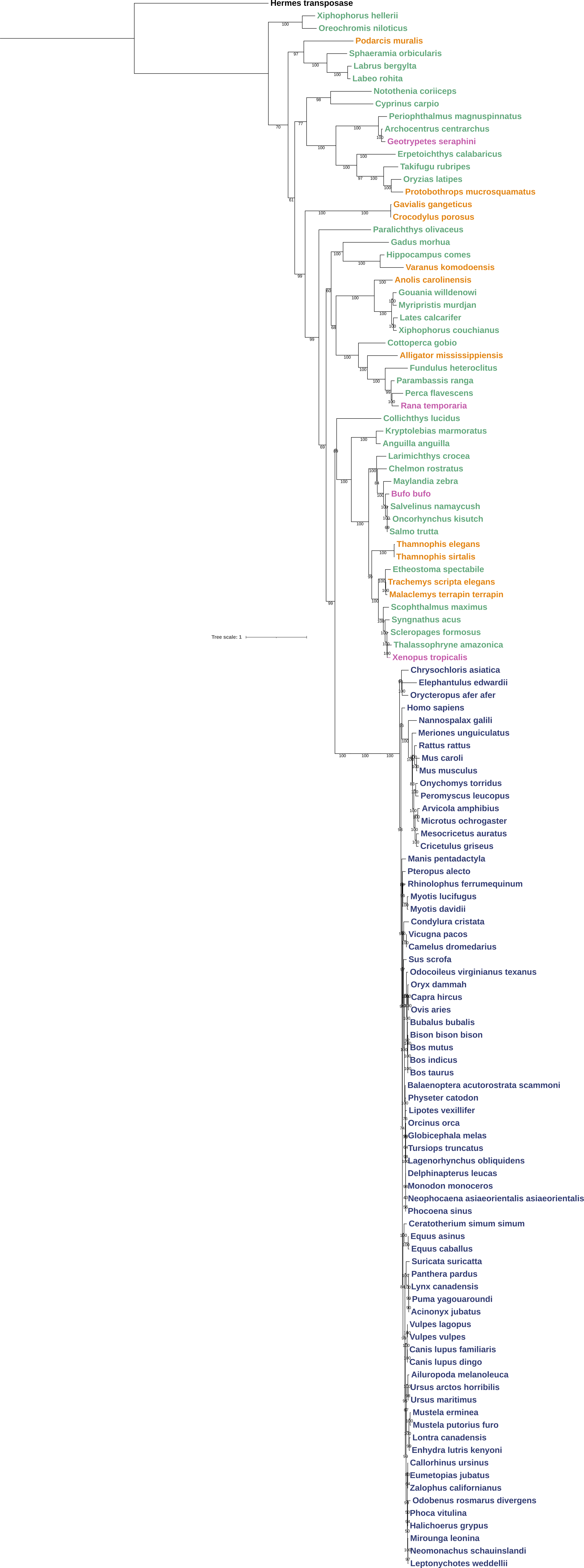
